# HighPlay: Cyclic Peptide Sequence Design Based on Reinforcement Learning and Protein Structure Prediction

**DOI:** 10.1101/2025.03.17.643626

**Authors:** Huitian Lin, Cheng Zhu, Tianfeng Shang, Ning Zhu, Kang Lin, Xiang Shao, Xudong Wang, Hongliang Duan

**Author notes:** The authors contribute equally to this work (Huitian Lin, Cheng Zhu). **Corresponding author**: **Hongliang Duan**: Macao Polytechnic University, Macao, 999078, China., **Xudong Wang**: College of Pharmaceutical Sciences, Zhejiang University of Technology, Hangzhou, 310014, P. R. China., **Xiang Shao**: Centralab, Suzhou Kowloon Hospital, Shanghai Jiao Tong University School of Medicine, Suzhou, 215028, P. R. China.

## Abstract

The structural diversity and good biocompatibility of cyclic peptides has led to their emergence as potential therapeutic agents. Traditional cyclic peptide design relies on natural template modification and combinatorial chemical library screening, but suffers from bottlenecks such as limited molecular diversity, high cost, and time-consuming.AI technologies have improved design efficiency by predicting target binding patterns and generating scaffold structures, but they still require extensive laboratory screening. In this study, we propose HighPlay, which integrates reinforcement learning (Monte Carlo Tree Search) with the HighFold structure prediction model to design cyclic peptide sequences for protein targets, dynamically exploring the sequence space without the need of predefined target information. The model was applied to the design of cyclic peptide sequences for three different targets, which were screened and verified by molecular dynamics simulation, and showed good binding affinity. Specifically, the cyclic peptide sequences designed for TEAD4 target showed micromolar-level affinity in further experimental validation.

## Introduction

The emergence of peptide drugs has filled the gap between small molecule chemical drugs and large molecule biologics and occupies an important place in modern drug development, as they are adept at binding to the substantial and often unstable surfaces on proteins that regulate PPIs.^1–3^ Cyclic peptides, a special class of peptide drugs, have a unique cyclic structure that promotes the formation of intramolecular hydrogen bonds and reduces molecular polarity, which in turn improves membrane permeability and makes it easier for them to penetrate cell membranes and reach intracellular targets.^4–6^ Meanwhile, the conformation of cyclic peptides is more rigid, which not only enhances their binding affinity and specificity to the target, but also reduces the likelihood of being degraded by proteases, thus improving their own stability.^7–10^ In practical applications, cyclic peptides have been widely used in various fields, including anti-infection, anti-tumor, and targeted therapies. They are employed in the treatment of conditions such as T-cell lymphoma^11^ and lupus nephritis.^12^

Traditional methods for cyclic peptide design rely on three main technological pathways: modification based on natural templates^13^, high-throughput screening of combinatorial chemical libraries^14,15^, and rational design based on structural biology.^16,17^ The natural template modification method improves the properties of known natural cyclic peptides (such as marine toxins^18^ or antimicrobial peptides^19^) by extracting their backbones for local amino acid substitutions, additions, or deletions. However, this method allows only limited modifications and struggles to break free from the constraints of natural structures, resulting in relatively low molecular diversity of the generated cyclic peptides. High-throughput screening of combinatorial chemical libraries requires the construction of compound libraries containing a large number of cyclic peptides with different structures^15^, For example, the design of peptides of 10 amino acids requires traversing 20^10^ (about 10^13^) possibilities, and is limited by synthetic methods, raw materials and other factors, making it difficult to achieve the true sense of complete randomness and infinite diversity of cyclic peptide combinations. The rational design oriented by structural biology relies on the three-dimensional structure of the target, active site and other information for cyclic peptide structure design, but it is not easy to obtain the three-dimensional structure information of the target by using X-ray crystallography^20^, nuclear magnetic resonance^21^ and other techniques, and some particularly complex targets lack a clear active site. These traditional methods have certain bottlenecks and require a large number of experimental procedures, which are costly and time-consuming to design.

In recent years, the rapid development of artificial intelligence technology has provided new ideas and methods for designing cyclic peptides. Among them, the emergence of structure prediction models such as AlphaFold2^22^, RoseTTAFold2^23^ has made it possible to predict the three-dimensional structure of proteins based on amino acid sequences. Deep learning has also been successfully applied to the de novo design of proteins and binding agents, for instance, using RFdiffusion^24^ to generate potential backbone structures and then ProteinMPNN to design sequences.^25,26^ However, this method requires thousands or even tens of thousands of design scenarios to be constructed during the computer simulation phase, which is extremely demanding on computational resources; it is difficult to achieve absolute precision in finding the sequence that best fits a given backbone structure; and extensive laboratory screening is required to identify an effective binding agent. EvoBind is a peptide design method based on AlphaFold, which aims to design peptide binders that are capable of binding to specific protein targets by means of computer simulations of directed evolution.^27^ However, this approach lacks effective guidance of the search space because it optimizes the peptide sequence by random mutation. In the complex protein-peptide interaction system, it is difficult to dynamically adjust the search direction based on previous search experience and feedback, and easily falls into the local optimal solution that cannot fully explore the large sequence space.

Reinforcement learning, as a technique that optimizes decision-making through interaction with the environment and based on reward mechanisms, can facilitate efficient exploration of sequence space. Reinforcement learning has demonstrated significant potential in solving combinatorial optimization problems across various fields, including chip placement design^28^, nuclear fusion control^29^, flow battery design^30^, matrix multiplication acceleration^31^, and autonomous driving.^32^ DeepMind pioneered the integration of Monte Carlo Tree Search^33^ (MCTS) with deep reinforcement learning, leading to the creation of AlphaGo^34^ and AlphaZero^35^, achieving mastery over multiple board games. EvoPlay applied reinforcement learning integrated with Monte Carlo Tree Search to design protein sequences, guiding exploration with both breadth and depth, and successfully designed a luciferase variant with a 7.8-fold improvement in bioluminescence compared to the wild type.^36^ Previously, our research group developed HighFold, a structure prediction model based on AlphaFold2, can precisely forecast the structures of cyclic peptides featuring head-to-tail cyclization and disulfide bonds, as well as their complexes, demonstrating exceptional predictive performance.^37^

Based on these principles, this study developed HighPlay, a cyclic peptide sequence design framework combining a reinforcement learning algorithm and the HighFold protein structure prediction model. The purpose of this combination is to dynamically explore the cyclic peptide space to generate suitable cyclic peptide binders based solely on the target protein sequence information. HighPlay employs Monte Carlo Tree Search (MCTS)-enhanced reinforcement learning algorithms to carry out residue mutation to optimize the sequence. Meanwhile, the architecture of the protein-cyclic peptide complex is predicted through HighFold, which identifies the suitable binding site and structure independently, without relying on any prior information. In this study, the framework was used in designing cyclic peptide binders targeting three different target proteins and the generated cyclic peptide binders were evaluated using molecular dynamics simulations, all of which demonstrated favorable binding affinities. The cyclic peptide binders designed to target the TEA domain family member 4 (TEAD4) was experimentally validated, successfully detecting micromolar-level binding affinity.

## Results and Disscusion

In this study, a novel cyclic peptide sequence design framework was developed, HighPlay, which integrates a reinforcement learning algorithm enhanced by Monte Carlo Tree Search (MCTS) with the deep learning-based HighFold protein structure prediction model. This framework is specifically designed for in silico directed evolution, enabling the design of cyclic peptide binders solely based on target protein sequence information. The framework employs a fully flexible sequence-structure co-evolution strategy: the initial cyclic peptide sequences were first generated by random initialization, followed by the HighFold model predicting the three-dimensional conformations and structural scores (with predicted local distance difference test (pLDDT)> 70 as the high-confidence criterion) of the protein-cyclic peptide complexes, and then a reinforcement learning strategy combining Transformer and MCTS guides the iterative mutation optimization of the cyclic peptide sequences. After preliminary evaluation using Rosetta Interface Analyzer, the final dominant sequences are selected and further validated through molecular dynamics simulations and surface plasmon resonance analysis (Figure 1A).

**Figure 1.**
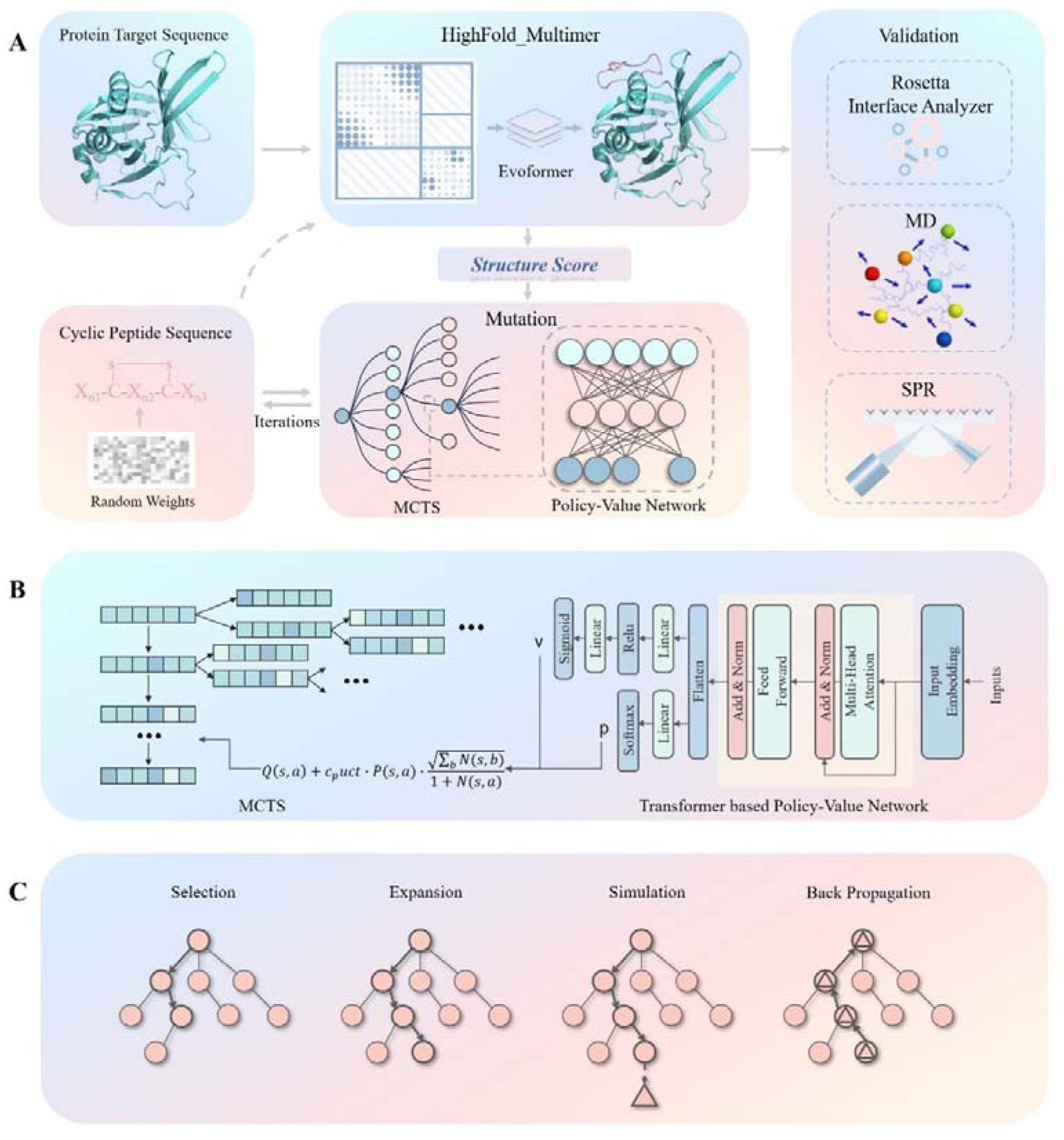
Design of cyclic peptide binders with HighPlay. (A) Schematic of the workflow of HighPlay. The input to the model is the sequence of the target protein. The initial cyclic peptide is randomly generated based on a probability distribution and consists of a disulfide bonded ring with two cysteines separated by a distance greater than 2/3 of the length of the sequence. The generated cyclic peptide is validated by Rosetta Interface Analyzer, Molecular Dynamics Simulation (MD), and Surface Plasmon Resonance (SPR) (B) Reinforcement learning strategy consisting of Monte Carlo Tree Search (MCTS) and Transformer-based policy-value network for iterative mutation of cyclic peptides. (C) The four steps performed for each iteration of Monte Carlo Tree Search.

HighPlay designs the cyclic peptides using policy-value network-guided MCTS, which is similar to playing a game on a chessboard. The preliminary sequence may be selected at random, or alternatively, the user may define an alternate sequence if they prefer. The optimization procedure of HighPlay comprises a series of consecutive play episodes. The state of the environment is depicted by means of a binary array of dimensions L×20, where L denotes the length of the cyclic peptide sequence and 20 signifies the number of amino acid types. In this context, a move refers to altering the type of amino acid at a position in the cyclic peptide sequence, that is, the distribution of a s single row in the state matrix. Within the context of each play episode, the reinforcement learning mechanism of HighPlay continues to perform mutation operations until the specified termination criteria are attained (Methods section). It is imperative to undertake multiple explorations using the Monte Carlo Tree Search prior to executing the mutation operation, including selection, expansion, simulation and back propagation (Figure 1C). During each exploration, the algorithm navigates through the Monte Carlo Search tree, beginning at the root node (representing the initial sequence state) and progressing until it reaches a leaf node. Each node within the tree is representative of a unique sequence. When the current node is in state *s*, the choice of the subsequent node is based on the largest sum of the values of Q(s,a) and U(s,a). These values are respectively decided by the relevant edges extending from the current node state (Methods section). When a leaf node with no child nodes, it is expanded and evaluated based on the outputs *p* and *v* from the policy-value network. The policy-value network, built on a Transformer^38^ architecture, evaluates the node state and predicts possible actions and their outcomes (Figure 1B).

HighPlay implements an iterative forward-looking MCTS process (exploration) to generate samples used to train a transformer-based policy-value neural network. This is subsequently employed in order to guide the search. In this process, the algorithm completely discards the preset constraints in traditional rational design (such as scaffold template specification or binding site limitation), enabling the system to autonomously explore uncharted chemical space and granting full autonomy in the generation of sequence-structure combinations. In traditional peptide binder design, prior knowledge and research experience often shape preconceived notions about ideal binding structures, sequences, or sites, which may deviate from actual conditions. By abandoning these potentially erroneous preconceptions, HighPlay ensures that the design process is not influenced by them, effectively avoiding ineffective exploration caused by prior bias and opening up a more objective and efficient pathway for the design of cyclic peptide binders. The designed binders were further validated using Rosetta Interface Analyzer, Molecular Dynamics simulations (MD), and Surface Plasmon Resonance experiments (SPR).

### Cyclic Peptide Binder Design

In this study, we employed HighPlay to design cyclic peptide binders for three different protein targets (PDB ID 1SSC、3R7G、6SEO).For each target, the cyclic peptide sequence length was set to 8-20 residues, and 13 designs were performed for each. After 300 iterations, cyclic peptide binders with pLDDT >90 were obtained. For the 1SSC target, eight out of the 13 designs achieved high-confidence cyclic peptide binders (pLDDT >70) after 300 iterations. Furthermore, it was observed that when the sequence length was set to 14, the highest pLDDT value of 90.15 was achieved.

The iterative optimization process of cyclic peptide binders is essentially a continuous extension and evolution of the Monte Carlo Tree (MCT). In the MCT tree structure, each node corresponds to a specific cyclic peptide sequence. By back propagate from the best cyclic peptide binder (leaf node), can sequentially find its parent node, grandparent node and so on up to the root node. The continuous branch formed by this series of nodes in the MCT search space can be referred to as the optimal path of the MCT (Figure 2A, B, C), illustrating the trajectory from the initial state to the optimal cyclic peptide binder state. To investigate whether the cyclic peptide binders generated during the Monte Carlo Tree search process in this study were optimized as expected, analyzed the pLDDT of the cyclic peptide binders along the optimal path of the MCT and calculated the corresponding protein-cyclic peptide complex scores (Score) using Rosetta Interface Analyzer.

**Figure 2.**
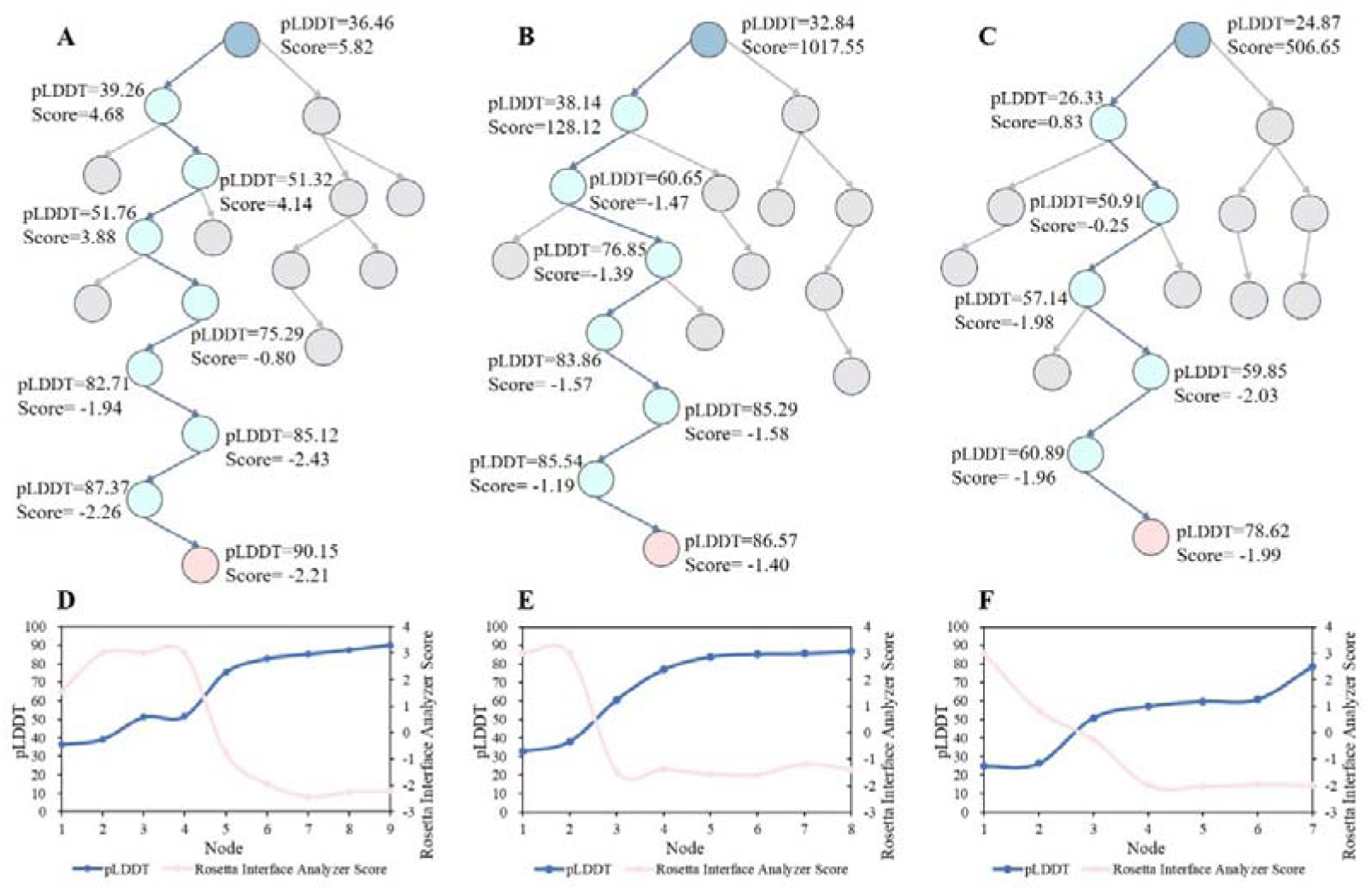
Optimal paths in Monte Carlo tree search space of the targets (A)1SSC (B)3R7G (C)6SEO. Changes in pLDDT and Rosetta Interface Analyzer Score of the corresponding cyclic peptides with node expansion on the optimal path branches of the targets (D)1SSC (E)3R7G (F)6SEO

Examining the changes of pLDDT and Rosetta Interface Analyzer Score of the nodes on the best path of the Monte Carlo tree, it was found that as the nodes in the Monte Carlo tree expanded, the pLDDT of the cyclic peptide binders continuously increased, and the corresponding Rosetta Interface Analyzer scores showed a roughly downward trend(Figure 2D, E, F).

### Molecular Dynamics Simulations

For each target, following the design of cyclic peptide sequences with lengths ranging from 8 to 20 residues, three dominant cyclic peptide sequence lengths were identified and the top 10 cyclic peptide binders were selected (Methods section). Subsequently, 100 ns molecular dynamics (MD) simulations were carried out for the designed protein-cyclic peptide complex structures. The MD simulation results indicated that the root-mean-square deviation (RMSD) of the protein backbones in the designed protein-cyclic peptide complexes converged to 3.0 Å (Figure 3), indicating that these cyclic peptide binders stably bind to their respective target proteins. Then, the binding energies of the protein-cyclic peptide complexes are determined by calculation using the method referred to as the molecular mechanics-generalized Born surface area (MM-GBSA) method. All the binding energies were found to be < –35 kcal/mol (Table 1), with the best binding energy recorded being –128.45 kcal/mol for the cyclic peptide binder designed for the target protein 3R7G.

**Figure 3.**
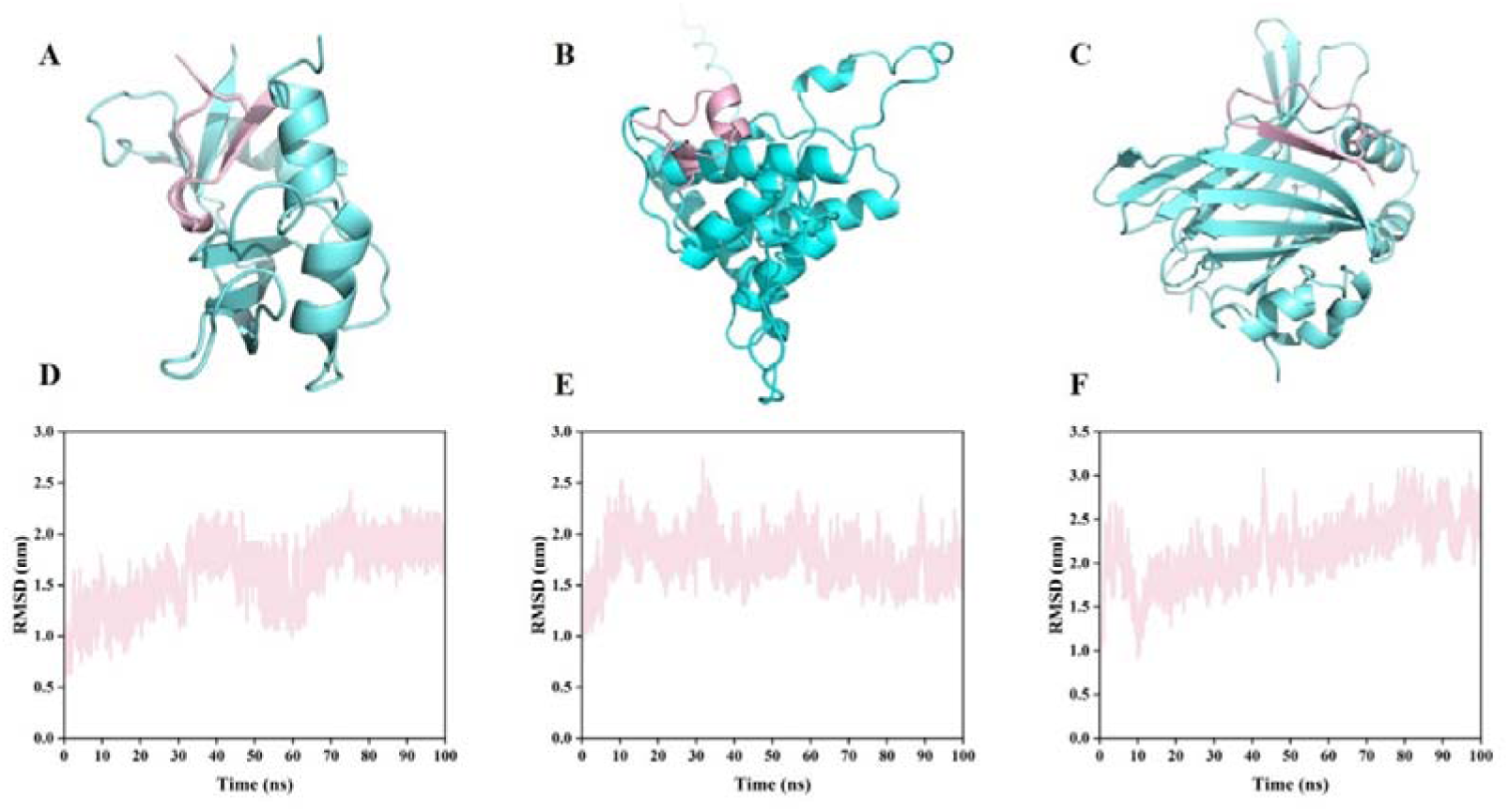
(A) Predicted structures of 1SSC and TPCYMENANDNDCK. (B) Predicted structures of 3R7G and SLCRCPVFQEYGLLTCT. (A) Predicted structures of 6SEO and PCQPMLARENTYMRCY. The predicted structures are shown with the target protein in cyan with the cyclic peptide binder in the form of pink stick. (D)The backbone RMSD of the binders along with the simulation time for A. (E)The backbone RMSD of the binders along with the simulation time for B. (F)The backbone RMSD of the binders along with the simulation time for C.

**Figure 4.**
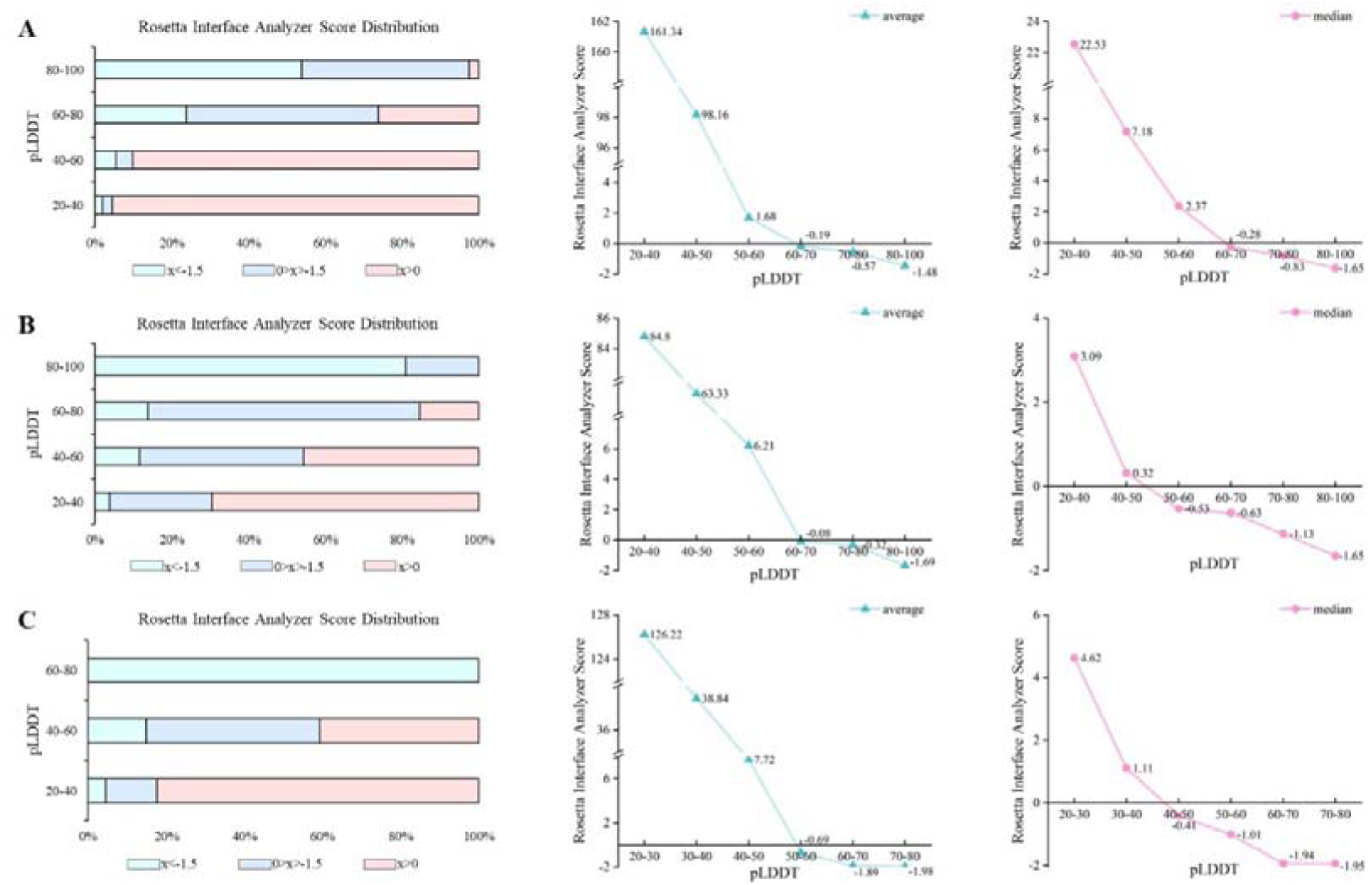
(A) Left: Distribution of Rosetta Interface Analyzer Score corresponding to different pLDDT. Center: Change in average Rosetta Interface Analyzer Score with increasing pLDDT. Right: Change in median Rosetta Interface Analyzer Score with increasing pLDDT. for target 1SSC. (B) Left: Distribution of Rosetta Interface Analyzer Score corresponding to different pLDDT. Center: Change in average Rosetta Interface Analyzer Score with increasing pLDDT. Right: Change in median Rosetta Interface Analyzer Score with increasing pLDDT. for target 3R7G. (C) Left: Distribution of Rosetta Interface Analyzer Score corresponding to different pLDDT. Center: Change in average Rosetta Interface Analyzer Score with increasing pLDDT. Right: Change in median Rosetta Interface Analyzer Score with increasing pLDDT. for target 6SEO.

**Table 1.**
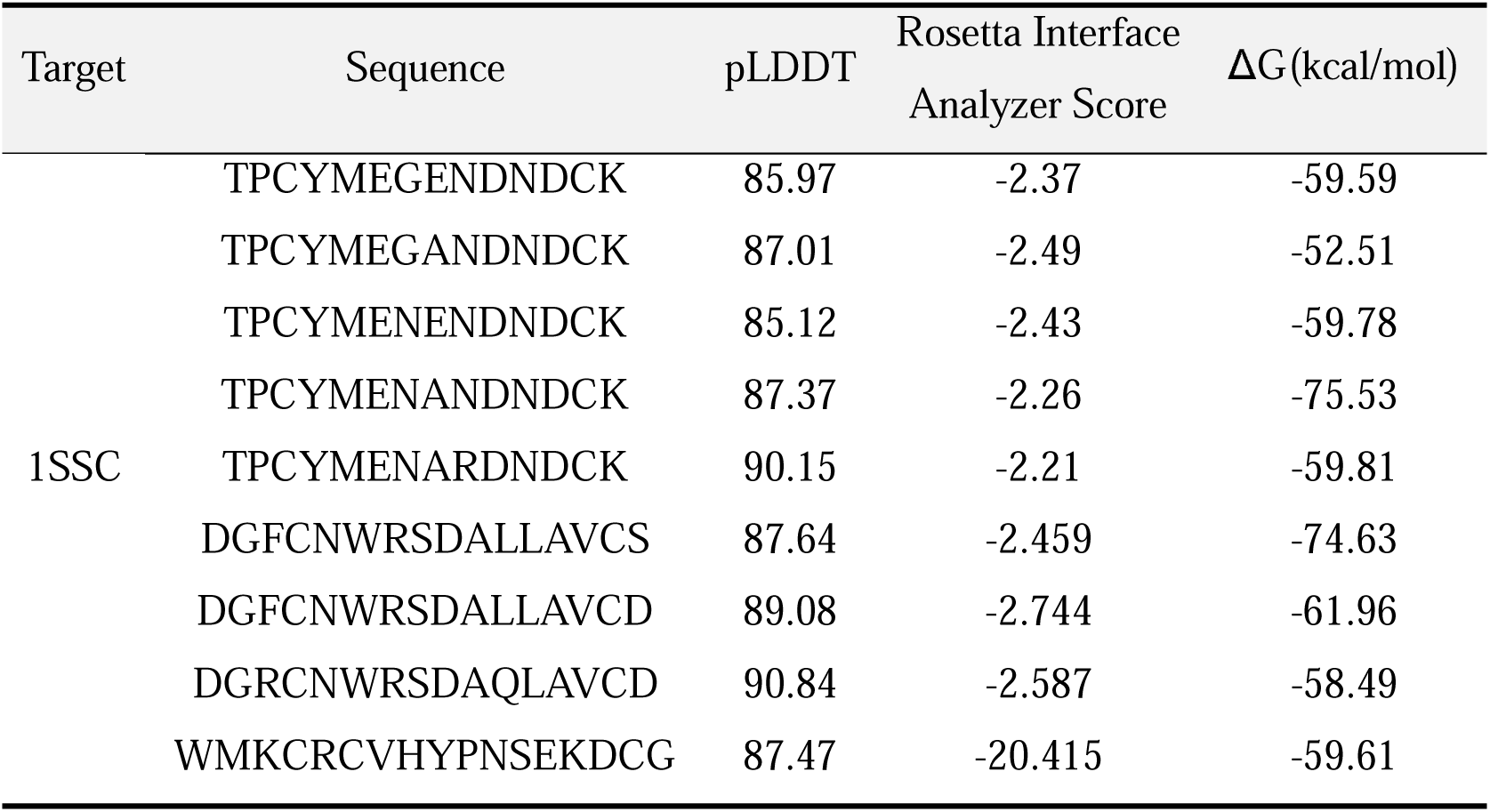
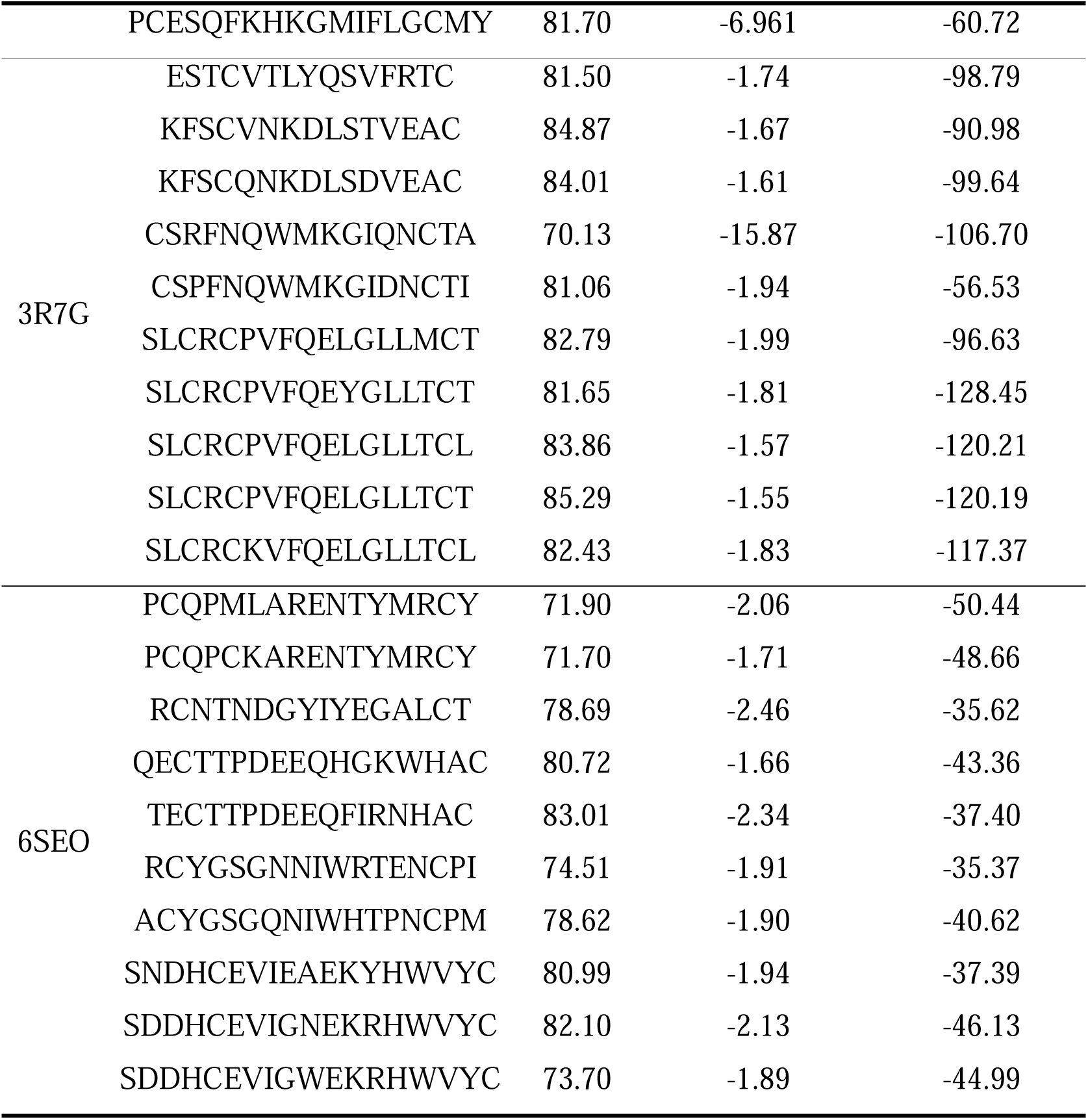
Cyclic peptide sequence analysis of target proteins 1SSC, 3R7G, 6SEO: pLDDT, Rosetta Interface Analyzer Score and binding energy (ΔG)

**Table 2.**
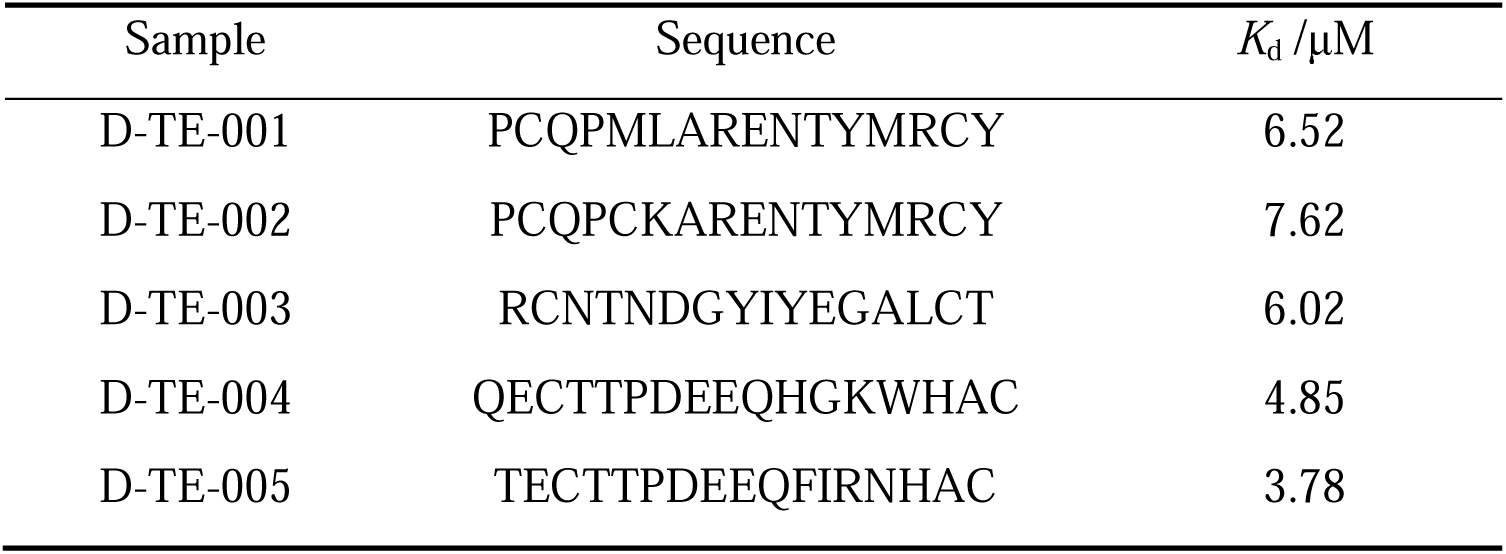
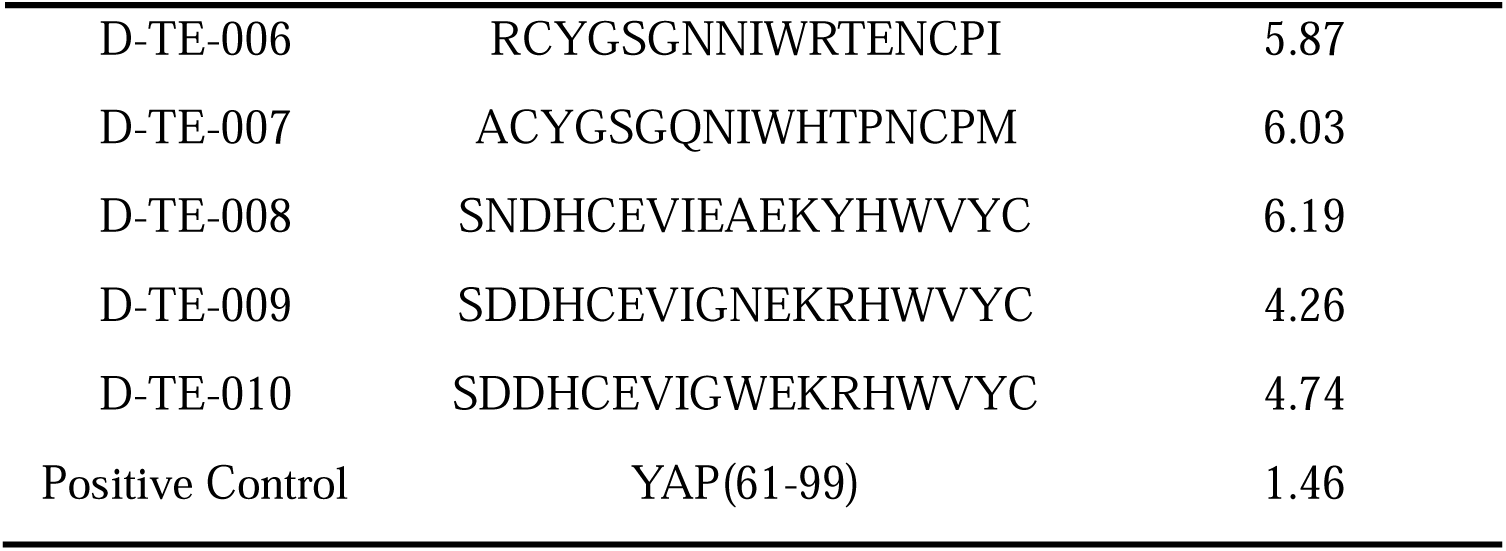
Binding affinity (*K*_d_) of TEAD4 to cyclic peptides and positive control.

### Surface Plasmon Resonance Experiments

The TEAD (Transcription Enhancement Associated Domain) proteins, acting as downstream effectors of the Hippo signaling pathway, play a critical role in regulating organ size and preventing tumor formation.^39^ TEAD4 (PDB ID 6SEO), an essential component of the TEAD protein family, regulates trophoblast differentiation during embryonic development^40^ and is involved in hematopoiesis by directly binding to the promoter region of the γ-globin gene to suppress fetal hemoglobin (HbF) expression^41^.In this study, the synthesis of the top 10 cyclic peptide binders designed for the TEAD4 target was conducted, and surface plasmon resonance (SPR) was used to measure the binding strength of these designed cyclic peptide binders to TEAD4. The purified TEAD4 protein was fixed onto a CM5 chip, and the cyclic peptides were employed as the analytes.

All 10 designed cyclic peptide binders all showed significant binding to TEAD4, and their corresponding dissociation constants (*K*_d_) calculated using a 1:1 Langmuir binding model, all falling within the micromolar range. Among them, D-TE-002 exhibited the best specific binding, with a *K*_d_ of 3.78 µM (Table 4).

### Correlation between Structure Score and Rosetta Interface Analyzer Score

In this study, the structural score pLDDT was used to generate high-affinity binders. To assess the capability of the structural score in differentiating genuine from false binders or forecasting affinity, an analysis was conducted of the relationship between the pLDDT of the peptide and the Rosetta Interface Analyzer Score. The pLDDT is a metric used to evaluate the confidence level of each residue in predicted protein structures, where elevated pLDDT scores signify greater confidence. Residues with pLDDT > 70 are considered confident. In this study, the structural score used was the pLDDT calculated for the residues of the cyclic peptide. Rosetta Interface Analyzer is an application within the Rosetta molecular modeling software suite, specifically designed to analyze protein-protein or protein-ligand interaction interfaces. It provides the dG_separated / dSASAx100, which assesses the energy efficiency of binding interfaces of different sizes. More negative values indicate more significant changes in binding energy per unit surface area, reflecting better binding stability and stronger binding affinity between the protein and the cyclic peptide. Typically, values below –1.5 are considered very good.

This study sought to analyze the correlation between pLDDT and Rosetta Interface Analyzer Score (dG_separated / dSASAx100) for three different targets. The results revealed that as the pLDDT values increased, the proportion of dG_separated / dSASAx100 < –1.5 also increased, while the proportion of dG_separated / dSASAx100 > 0 gradually decreased or even reached zero. By grouping all cyclic peptide binders based on their pLDDT values and calculating the mean and median Scores, we observed that as pLDDT increased, the Scores consistently showed a decreasing trend. This indicates a negative correlation between pLDDT and Rosetta Interface Analyzer Score to some extent, suggesting that pLDDT can serve as an effective indicator for predicting binding affinity.

## Conclusion

The use of artificial intelligence to design proteins and peptide binders presents a significant opportunity to expedite the expansion of protein diversity available for biotechnological applications. HighPlay developed in this study allows the design of suitable cyclic peptide binders for target proteins when only the sequence information of the target protein is available. This method doesn’t require prior information about binding sites, template structures, and binder dimensions, rendering it highly adaptable to various targets.

HighPlay applies reinforcement learning to the design of cyclic peptide binders, using a policy-value network based on Transformer to guide Monte Carlo Tree Search (MCTS). Through intelligent decision making and dynamic optimization, this reinforcement learning approach significantly improves the efficiency of cyclic peptide design. In the complex environment of cyclic peptide design, the reinforcement learning agent treats the cyclic peptide sequence as the action space and continuously interacts with the environment (protein target). Guided by reward signals (structural scores), the agent rapidly learns and adjusts its strategy. This method avoids the blind search inherent in traditional approaches and, through accumulated experience, precisely focuses on exploring high-potential regions of cyclic peptide sequences, effectively reducing futile attempts. Moreover, during the iterative process, the agent can quickly optimize sequences based on real-time feedback, enabling the acquisition of high-performance cyclic peptide designs in a shorter time frame. This significantly improves design efficiency and provides robust support for applications such as cyclic peptide drug development.

In this study, we conducted affinity measurements for the cyclic peptide binders designed to target the TEAD4 protein. In order to validate the functionality of the expressed protein for the aforementioned affinity measurements, we synthesized and measured the affinity of another protein known as YAP(61-99). This protein was used as a positive control, and the resulting affinity value was determined to be 1.46 µM. The results showed that although the 10 designed cyclic binders did not outperform YAP(61-99) in terms of affinity, they all exhibited binding affinities at the µM level. This demonstrates that HighPlay is capable of efficiently designing cyclic peptide binders that exhibit strong affinity for the target protein, demonstrating its potential to develop novel high-affinity cyclic peptide binders. It is essential to be able to explore new protein targets without the need to predefine specific binding sites or limit the size of binders. This method offers significant promise for expanding the number of proteins available in various biotechnological applications.

## Methods

### HighPlay Framework

The environmental state of HighPlay is depicted by an L×20 binary matrix, where the rows represent positions (L) and the columns represent amino acids (20 types). The corresponding action space is a flattened vector with a size L×20. Upon performing an action, an object is selected from the action space vector, resulting in a single point mutation within the state sequence. This mutation has a clear representation in the state matrix: the object specified by the action is assigned a value of 1, whereas the other objects, which correspond to different residue types within the same row (corresponding to a specific position in the peptide), are marked as 0.An episode comprises a sequence of actions, with associated states represented as s_i_ (0 < i< N) and the rewards indicated as r_i_(0 < i < N). State update is an iterative process that continues until meets its termination criteria. The first end criterion is that the reward score r_i_ of the present state is less than the reward r_i-1_ of the previous state, indicating that the execution of the action failed to result in an increase in reward. The other criterion is that the sequence of states after the execution of the action transforms into a sequence that has already occurred before, which is called action invalidation.

It is evident that the tree search algorithm employed by HighPlay is comprised of four distinct phases of MCTS, selection, expansion, simulation, and back propagation, respectively. The MCTS exploration process is guided by a policy-value neural network. Each exploration is associated with an episode and traverses from the root node, designated as s_root_, to the leaf node, denoted as s_L_, within the Monte Carlo tree. It is important to note that each node in this structure corresponds to s a distinct state, or sequence, within the context of the exploration. To move from the current state s to the next state s’, an action a need to be performed. The edge connecting s and s’ is labeled as (s, a). During exploration, the selection of subsequent nodes is determined by the maximum value of the sum of Q(s,a) and U(s,a). Here, Q(s,a) represents the action value associated with the edge (s,a), which is defined as the expected payoff from executing action a when in state s. The evaluation and subsequent update of this value is conducted by the output v from the neural network, along with the visits count N(s, a). U(s,a) is an exploration term that balances exploration and exploitation. It is determined through the following equation:

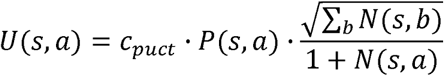

where c_puct_ is a constant that balances exploration and exploitation. P(s,a) denotes the prior probability of moving from parent node s to a child node. This probability is determined by inputting the state of the parent node into the neural network. Upon reaching a leaf node s_L_, the search algorithm expands the tree by generating new child nodes. Finally, the tree is updated backwards, proceeding from the leaf nodes towards the root node.

When simulation of the root node(s_i_)is complete, an action r_i_ is sampled based on a specific probability distribution.

Specifically, the action is proportional to the access count raised to a power at time step i, the mathematical relationship can be expressed as follows 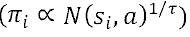, where τ is an adjustable temperature parameter. The reward r_i_ for every action is assessed through an agent model. The data generated from each episode (*s_i_, r_i_, r_i_*) is stored in a dedicated buffer for reinforcement learning. This buffer functions as a data repository, continuously accumulating information from various episodes, thereby providing a rich sample resource for subsequent neural network training. The neural network(*p, v*) = *t_0_*(*s*) is trained with a combined loss term:

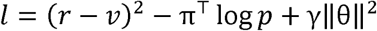

where *r* is the actual reward, *v* and *p* are the predicted state value and mutation probability, respectively, computed by the Transformer-based policy-value network, and γ is the regularization parameter for e. The episode terminates if the reward for the present action is less than the reward of the previous action, and if no improvement in adaptation is observed for thirty consecutive episodes, a new starting sequence can be chosen from the pool of sequences to begin a new episode.

### Transformer-based Policy-Value Networks

The policy-value neural network receives as input a state matrix of dimensions L×20, where L signifies to the length of the cyclic peptide and 20 signifies the total number of amino acid types. This model architecture based on Transformer is composed of an embedding layer, a Transformer encoder, and two independent output heads (policy head and value head) that process the input state matrix. This architecture not only learns complex patterns and dependencies but also simultaneously evaluates the value of the state and guides the selection of the next action, making it well-suited for tasks requiring fine-grained feature extraction and decision-making.

### Rosetta Interface Analyzer

In this study, the Rosetta interface analysis tool was used to precisely compute the Rosetta interface score (dG_separated/dSASA × 100) of the generated protein-cyclic peptide structures.^42^ The procedure was as follows: first, based on the initially predicted complex structure, the “minimize” function was used to search for and obtain the structure with the lowest energy. Considering that there is a possibility that the cyclic peptides in the complex can be converted into linear peptides, to avoid this situation, “-use_truncated_termini” and “-relax:bb_move false” were specifically enabled during the operation. Subsequently, the minimized complexes were used as input data for the “InterfaceAnalyzer”. When calculating the binding energy, the “-pack_separated” option was used in order to rearrange the exposed interfaces. In addition, all Rosetta applications involved in this study used the standard weights REF15 and the running environment was Rosetta 3.13 Linux version to ensure the consistency of the experimental conditions and the reliability of the results.

### Selection of Designed Cyclic Peptide

For each target, cyclic peptide designs were carried out with sequence lengths between 8 and 20 residues. The proportion of cyclic peptides with pLDDT >70 was calculated for each peptide lengths, and the top three sequence lengths with the highest proportions were selected as the dominant lengths. The cyclic peptide sequences generated at these dominant lengths were evaluated using Rosetta Interface Analyzer, and those meeting the criteria of pLDDT >70 and dG_separated/dSASA×100 < –1.5 were selected. From these, a total of 10 cyclic peptide sequences were chosen for further validation.

### Molecular Dynamic Simulation

Molecular dynamics simulations were performed using the pmemd.cuda module in Amber 24 for 100 ns. Peptides and proteins were parameterized using the ff19SB force field.^43^ The protonation states of the residues were preconditioned using pdb2pqr.^44^ The simulation system was solvated in a truncated octahedral box filled with OPC water molecules, with the box edge 10 Å away from the system, and sodium or chloride ions were added to neutralize the charge of the system. Long-range electrostatic interactions were treated using the particle mesh Ewald (PME) method with a cutoff distance of 8 Å. The system underwent a stepwise equilibration process: first, a harmonic restraint of 2.0 kcal/(mol·Å²) was imposed on the protein and peptide to avoid excessive deformation, and energy minimization was performed with 500 steps of steepest descent and 500 steps of conjugate gradient. After energy minimization, the protein and peptide were kept under a 2.0 kcal/(mol·Å²) restraint to prevent violent motion and heated from 0 K to 300 K for 25 ps under NVT conditions with Langevin temperature control. Subsequently, a 50 ps NPT simulation was performed at 1 atm pressure with a 2 kcal/(mol·Å²) harmonic restraint on the complex. Then, a 500 ps unrestrained equilibrium simulation was performed in the NPT ensemble, with the temperature and pressure controlled by the Langevin temperature controller and the Berendsen pressure controller, respectively. Finally, a 100 ns unrestrained production simulation was performed in the NPT ensemble, with the temperature and pressure maintained by the Langevin thermostat and the Berendsen pressure controller, respectively. The simulation time step was 2 fs, and the SHAKE algorithm was used to constrain hydrogen atoms. Finally, the root mean square deviation (RMSD) in the trajectory was analyzed using the CPPTRAJ module^45^, and the binding free energy of the protein and peptide was calculated using MMPBSA.py.^46^

### Synthesis of Cyclic Peptides

The cyclic peptides designed by HighPlay were synthesized by means of standard Fmoc solid phase peptide synthesis (SPPS). 0.5 g of Rink amide MBHA resin was suspended in a freshly prepared solution consisting of 0.3 mmol of Fmoc-Cys(Trt)-OH, 0.3 mmol of HOBT dissolved in 8 ml of DMF, and 0.5 ml of DIC. The reaction was carried out under nitrogen bubbling conditions for 1.5 hours. After, the resin was filtered and washed at least three times with DMF and DCM, the Fmoc protecting groups were removed using a 20% piperidine/DMF solution. Subsequent amino acid coupling reactions were carried out by adding a freshly prepared solution (containing 0.9 mmol Fmoc-AA-OH, 0.9 mmol HOBT and 1 ml DIC) to 10 ml DMF. The reaction is carried out under nitrogen bubbling conditions for 1 hour. The repetition of deprotection and coupling steps leads to the formation of peptides of the desired length.

The peptide was cleaved from the resin using a cleavage cocktail of TFA/TIS/HLO (95:2.5:2.5) and precipitated with cold diethyl ether (EtLO). The precipitated peptide was redissolved in DMSO. The pH of the solution was adjusted to 8.0 and incubated for 2 h at room temperature under air oxidation to promote the formation of disulfide bonds. Finally, the pH of the solution was adjusted to 3-4 with trifluoroacetic acid (TFA) and purified by reversed-phase high performance liquid chromatography (RP-HPLC). The quality of the synthesized cyclic peptides was verified by high performance liquid chromatography (HPLC) and mass spectrometry (MS), and the purity was above 95%. The HPLC chromatograms and mass spectra of all compounds have been included in the Supporting Information.

### Affinity Measurement Using Surface Plasmon Resonance

In order to assess the binding affinity of the cyclic peptides for TEAD4, surface plasmon resonance (SPR) was employed. This technique is an optical-based, label-free detection method that allows for precise measurements.^47^ It is used to detect in real – time the binding interaction by detecting refractive – index changes near the sensor surface, which occur due to interactions between molecules on the sensor surface and their binding partners in the solution. The surface plasmon resonance experiments were conducted with a Biacore T200 device (GE Healthcare) using a Series S Sensor Chip CM5 (Cytiva). We immobilized the TEAD4 protein on the active flow cell using amine coupling of the protein to the surface of the chip. For this, standard amine coupling reagents 1-ethyl-3-(3-(dimethylamino)propyl)carbodiimide hydrochloride (EDC) and N-hydroxysuccinimide (NHS) were used to activate the sensor chip, then TEAD4 protein (at a concentration of 50 μg/mL, in 10 mM sodium acetate buffer, pH 4.0) was injected into the chip at a rate of 10 μL/min for coupling. Subsequently, the flow cell was blocked with 1 M ethanolamine to make the immobilization level of the TEAD4 protein be around 7000 resonance units (RU). The interaction experiments occurred at 25 °C within a running buffer consisting of 10 mM PBS (pH 7.4), 0.05% Tween 20, and 0.5% DMSO. The compounds were diluted to 10 μM in the running buffer and injected over the immobilized target protein. The flow rate was established at 30 μL/min, with an association time of 120 seconds, and a dissociation time of 120 seconds. In addition, cyclic peptides within different concentration ranges were used as the analytes. The highest concentration was set at 20 μM, and then ten concentration gradients were prepared through two-fold serial dilutions.

The data was processed and analyzed with the aid of Biacore Insight Evaluation Software. The raw sensorgram data were then subjected to a preprocessing procedure that entailed the subtraction of the response of the reference channel, in addition to the execution of corrections for blank injections. The sensor profiles were then fitted to a 1:1 binding model using two global kinetic parameters, the association rate constant (*k*_a_) and the dissociation rate constant (*k*_d_), as well as the maximum response value (R_max_) for each sample. The data retained during the fitting process were used to accurately determine the association rate constant (*k*_a_), the dissociation rate constant (*k*_d_), and the dissociation constant (*K*_d_). where the equilibrium dissociation constant (*K*_d_) is determined by the ratio of the dissociation rate constant (*k*_d_) to the binding rate constant (*k*_a_).

## Author Contributions

H.D. and X.W. conceived and supervised the research project. H.L., C.Z., N.Z., K.L. performed the dry experiments and analyzed the data; X.S. performed the wet experiments and analyzed the data; H.L., T.S. and N.Z. wrote the manuscript. All authors have read and approved the final version of the manuscript for submission to this journal.

## Acknowledgements

This project was supported by the Natural Science Foundation of Zhejiang Province (LD22H300004), the Pre-research Project of Suzhou Kowloon Hospital (SZJL202203) and the Applied Basic Research of Suzhou Medical and Health Science and Technology Innovation(SKY2023113).

